# Separable Codes for Time and Decision in Human Temporal Perception

**DOI:** 10.64898/2026.04.28.721130

**Authors:** Mateus Silvestrin, Peter Maurice Erna Claessens, Nicholas Myers, Andre Mascioli Cravo

## Abstract

Time perception involves estimating physical durations and making categorical judgments relative to reference intervals. However, most studies conflate these processes, limiting insight into how they are encoded in brain activity. Here, we used EEG and multivariate pattern analysis (MVPA) to dissociate neural representations of time and decision during a temporal discrimination task. Thirty participants (humans of any sex) compared variable intervals to block-specific references, with duration and category (shorter, equal, or longer) manipulated orthogonally. Behaviorally, responses were shaped by target duration, category and recent trial history. An Internal Reference Model (IRM) indicated that participants dynamically updated their internal reference over trials. MVPA showed that both physical duration and categorical decision information were encoded throughout the trial, though with distinct temporal profiles. These signals were represented along separable neural dimensions, enabling their differentiation in brain activity. These findings suggest that time perception relies on parallel, functionally distinct processes for tracking duration and making temporal decisions, supporting models that treat them as independent components of temporal evaluation.

## Introduction

Humans and non-human animals use time to interact with their environment (Merchant et al., 2013; Paton & Buonomano, 2018). Among temporal abilities, interval timing, the ability to estimate the duration of an interval, is one of the most studied. Models developed to explain how organisms track time can be classified in various ways, such as whether they rely on adaptive threshold settings or adaptive accumulation rates (Balcı & Simen, 2016). For example, Scalar Expectancy Theory (SET, Gibbon, 1977) proposes that organisms measure time by accumulating pulses from a pacemaker at a constant rate. In temporal categorization tasks, in which participants compare an interval to a learned reference, the accumulated count is evaluated against a reference memory. SET represents different durations by varying the number of accumulated pulses, consistent with an adaptive threshold mechanism.

In contrast, the Time-Adaptive Opponent Poisson Drift-Diffusion Model (TopDDM) combines diffusion processes to model both timing and decision-making (Balci & Simen, 2014; Simen et al., 2011). For temporal categorization, TopDDM employs a two-stage sequential diffusion process: first, accumulating noisy clock signals to time an interval, then accumulating evidence from that first stage to reach a decision. This model adjusts the rate of evidence accumulation in the first stage to reflect the expected duration, aligning with an adaptive rate framework.

Although these models rely on distinct principles, they often produce similar behavioral predictions (Balcı & Simen, 2016). SET and TopDDM are not unique in this respect, and the threshold-versus-rate distinction cuts across the broader landscape of timing models, yet this algorithmic difference is hard to see at the behavioral level. One possibility is to examine the neural correlates of timing. Threshold-based models predict a similar buildup of neural activity across intervals to a different threshold. In contrast, rate-based models predict rescaled activity buildup to the same level. However, characterizing duration and decision-related activity is a necessary step to adjudicate between models, as apparent rescaling may reflect changes in the decision variable rather than in the timing mechanism.

In EEG research, studies have investigated temporal processing by examining event-related potential (ERP) components such as the contingent negative variation (CNV), P300, and the late positive component (LPC). There is no clear consensus on what these ERPs indicate (Kononowicz et al., 2016), or whether brain activity before or after a timed interval is the most reliable marker (Bueno & Cravo, 2021; Kononowicz & van Rijn, 2014; Kruijne et al., 2020; Ofir & Landau, 2022). Most EEG studies use a small set of time intervals and a single reference, meaning that physical duration and categorical status are strongly correlated (i.e., longer intervals are always ‘longer than reference’ and shorter intervals always ‘shorter than reference’), making it impossible to determine whether brain activity reflects duration encoding, categorical decision formation, or both. Most studies also treat the learned references as static, although humans continuously update their internal representations of duration (Dyjas et al., 2012; Wiener et al., 2014). Failing to account for this recalibration risks attributing learning-related neural variance to timing mechanisms.

In this study, we address these limitations by investigating temporal discrimination while separating the contributions of physical duration and categorical judgment. We designed a task in which physical durations and their category (shorter, equal, or longer than a reference) varied orthogonally, allowing us to test whether duration and category are encoded by shared or separate neural mechanisms. We find support for separable encoding: duration information is most strongly represented before interval offset, while categorical decision signals peak afterward, each carried along separate neural dimensions. We also find that participants dynamically update their temporal references within and across blocks, and that these shifts are reflected in the neural signals underlying categorical judgments. These results speak to both major classes of timing models and also highlight the difficulty of disentangling changes in timing mechanisms from changes in the decision processes that operate on their outputs.

## Materials and Methods

### Data availability

All materials resulting from this study are openly available. Please see https://zenodo.org/records/22660211 for the raw and eeg files and https://zenodo.org/records/22660752 to access the task and analysis code.

### Participants

Thirty-six students (18 men) participated in the experiment. The sample size was based on similar EEG experiments (Damsma et al., 2021; Ofir & Landau, 2022). The average age of participants was 21.5 ± 2.8 years (mean ± sd). All participants were informed about the procedures and signed an informed consent form before the experiment, which was approved by the UFABC Research Ethics Committee. One participant performed at chance level on the task, likely due to limited attention or understanding. The electrophysiological signals from five other participants were too noisy (more than 30% of epochs were rejected due to noise). All analyses were conducted on the remaining 30 (15 men) participants who met both the behavioral performance criterion (above chance level) and the EEG data quality criterion (fewer than 30% of epochs rejected).

### Stimuli and procedure

The experiment consisted of a temporal discrimination task (Fig. 1a) combined with EEG recordings. The stimuli were presented using PsychoPy (version v3.0.0b11) (Peirce et al., 2019) on a ViewPixx monitor with a vertical refresh rate of 120 Hz, positioned approximately 52 cm from the participant. Responses were collected via a response box with nine buttons (DirectIN High-Speed Button; Empirisoft). Participants used the index fingers of both hands to respond with the extreme left and extreme right buttons.

**Figure 1.**
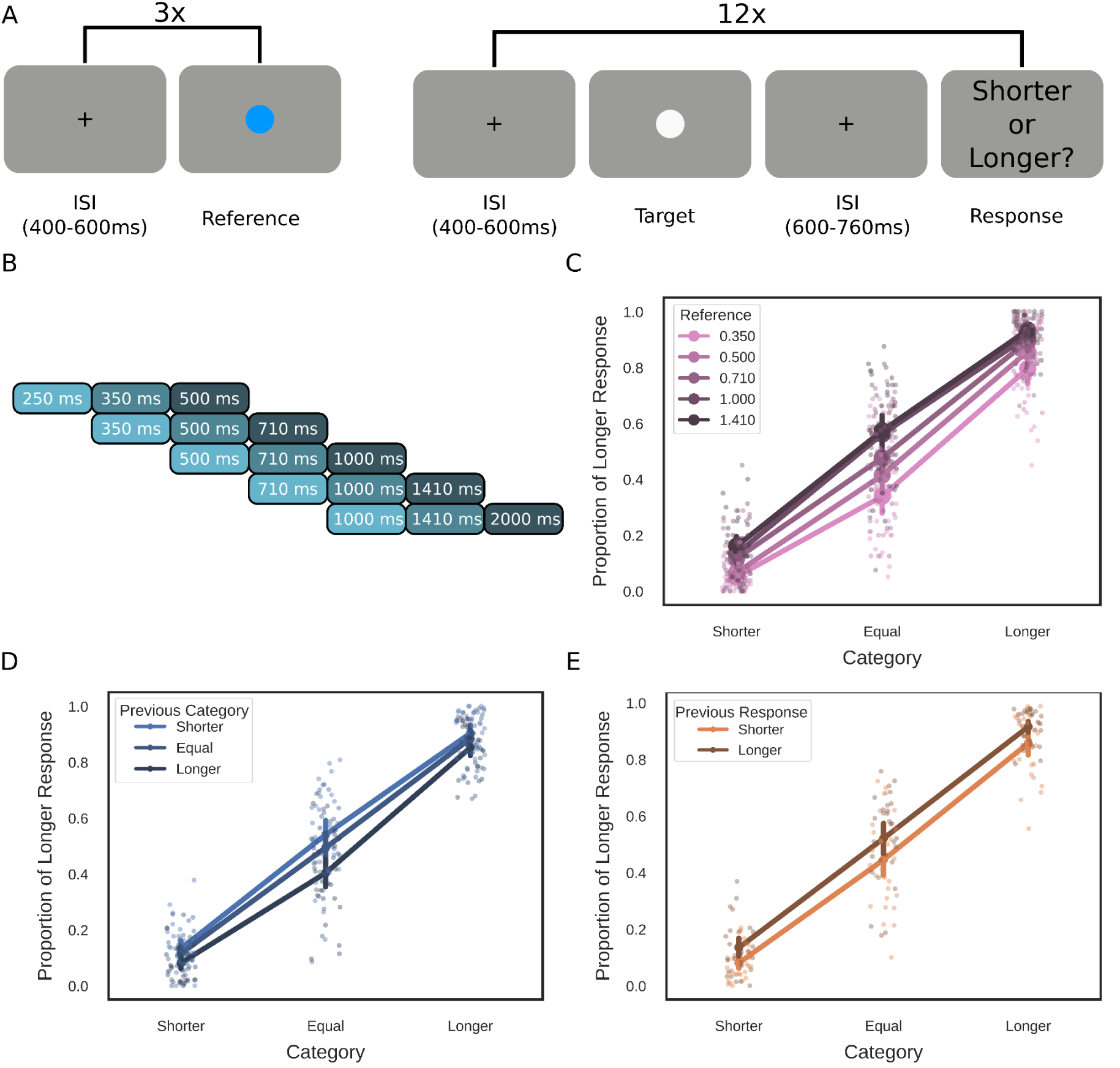
Experimental design and behavioral results. (A) Trial structure showing the temporal discrimination task. At the beginning of each block, participants viewed a reference interval (blue circle) presented three times. In each trial, a target interval was presented, and participants then made a categorical judgment (shorter or longer than the reference). (B) Duration scheme across experimental blocks. Logarithmically spaced intervals (250-2000 ms) were used, with different durations serving as references (350, 500, 710, 1000, 1410 ms) across blocks. Note that three durations (500, 710, 1000 ms) occupied different categorical positions (shorter, equal, longer) depending on the reference. Shades of blue indicate categorical position. (C) Behavioral performance showing proportion of “longer” responses as a function of target duration, separated by categorical status (shorter, equal, longer than reference). (D) Influence of previous trial category on current responses. (E) Influence of prior responses on current responses, indicating response repetition bias. Individual participant data are shown as light points, with group means and error bars representing 95% confidence intervals.

The experiment was designed so that the targets’ physical duration and category (shorter, equal, or longer than the reference) were orthogonal for a given set of durations. Each block began with the presentation of the reference for that block, represented by a blue circle in the center of the screen. The reference was presented three times, with an interstimulus interval (ISI) of 400-600 ms. During the block, the target durations were presented as white circles at the center of the screen. The mini-block structure, in which the reference was presented at the start of each block rather than on every trial, was chosen because it introduces trial-by-trial variability in the memorized interval, which can be modeled and linked to EEG activity. Following the duration offset, a response screen was presented after an inter-stimulus interval (ISI) of 600-760 ms. On this screen, volunteers indicated, by pressing a button on a response box, whether the interval was shorter (left button) or longer (right button) than the reference interval. Each block consisted of 12 trials, with four trials in each category (shorter, equal, or longer than the reference), presented in random order. The experiment consisted of 100 blocks and lasted approximately 90 minutes.

A diagram of the durations used in different blocks is shown in Figure 1b. Each block was organized around a central reference duration (350, 500, 710, 1000, or 1410 ms), with three possible intervals within each block: shorter, equal to, or longer than the reference. All intervals used in the experiment were logarithmically spaced. This design ensured that three durations (500, 710, and 1000 ms) were assigned to distinct categories (shorter, equal, or longer) across blocks. Participants were informed that some targets could be very close to the block’s reference duration and were instructed to respond as faithfully as possible to their perception in these cases of maximum uncertainty, rather than adopting a strategy of always selecting the same response (e.g., always “longer“) whenever in doubt. At the end of each block, participants received general feedback on the percentage of correct answers in that block. Responses to durations identical to those in the references were not included in the feedback as there were no objective correct answers. The equal trials were excluded solely from the feedback calculation, but were retained in all further analyses.

### Behavioral analysis

For the behavioral analysis, we used a logistic regression to estimate the influence of parametric quantities, such as target duration, category (shorter, equal, longer), previous category, response, and previous response, on the probability that participants responded longer than the reference. All continuous predictors were standardized to z-scores before entering the logistic regression models. Regressions were performed at the participant level, and the estimated coefficients were tested at the group level using standard parametric tests (e.g., t-tests and repeated-measures ANOVA). Unless otherwise stated, values in the text are reported as mean ± s.e.m.; error bars in figures represent 95% confidence intervals

### Internal Reference Model

To model whether and how participants dynamically update their internal references, we used a framework based on the Internal Reference Model (IRM) (Dyjas et al., 2012). In this model, participants compare the internal representation of X_n_ (the interval presented in the current trial) with an internal reference i_n_ (Dyjas et al., 2012). This internal reference is updated at the end of every trial, based on the presented interval at that trial, according to:

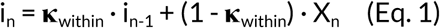

where i_n_ is the new updated internal reference at the end of trial n, i_n-1_ is the internal reference from the previous trial, and X_n_ is the duration presented in the current trial. Parameter **κ**_within_ controls the retention of the internal reference across trials, ranging from 0 (no retention/full updating, in which participants use the previous interval as the reference for each trial) to 1 (perfect retention/no updating, with perfect memory of the correct reference). When the IRM was adapted to a single-stimulus paradigm (in which a single interval is presented on each trial, as in our experiment), a single temporal reference was used throughout the experiment (i.e., participants learned the temporal reference at the start of the experiment and used the same reference throughout the whole task). However, our experimental task consisted of multiple short blocks (100 in total), each with 12 trials, and each began with three presentations of the reference interval. Thus, participants had to explicitly learn a new temporal reference in each block. To assess whether participants could learn the new reference fully or whether the internal reference used in the last block also modulated performance, we implemented a second update that accounts for between-block updating of the reference. In this between-block updating, at the start of each block, the internal reference was updated based on the three reference presentations according to:

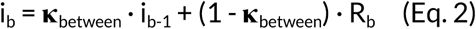

where R_b_ is the reference duration for the current block, and i_b-1_ is the latest internal reference. For the first block, the internal reference was initialized to the presented reference duration, as no prior internal reference existed. From the second block onward, the between-block update (Eq. 2) was applied using the internal reference carried over from the end of the previous block. This update was applied three times in succession to simulate the effects of the three reference presentations. Parameter **κ**_between_ controls the retention of the internal reference across blocks, with **κ**_between_ = 0 indicating full updating (complete reset to the new reference) and **κ**_between_ = 1 indicating complete retention of the previous internal reference (ignoring the reference presentations).

Within this framework, a perfect observer would have **κ**_within_ = 1 and **κ**_between_ = 0: throughout the block, it maintains the learned reference in memory without interference from the presented comparison intervals (perfect retention, **κ**_within_ = 1), and at the beginning of each new block, it fully updates its internal reference to the new reference with no carry-over from the previous block (complete updating, **κ**_between_ = 0). Note that although both parameters reflect retention, they do so with respect to different quantities: **κ**_within_ reflects resistance to unwanted interference from comparison intervals, while **κ**_between_ reflects resistance to updating from new reference presentations. The direction of desirable behavior therefore differs between them.

To estimate **κ**_within_ and **κ**_between_ for each participant, we performed a two-dimensional grid search, with both parameters varying independently from 0 to 1 in steps of 0.02 (51 × 51 = 2,601 parameter combinations per participant). For each parameter combination, we estimated the internal reference in each trial using Equations 1 and 2 and computed the distance from the internal reference as log(target_duration/ internal_reference). A logistic regression model was then fitted to predict participants’ responses, using this distance as the predictor variable and an intercept. The log-likelihood was computed for each parameter combination, and the combination yielding the highest log-likelihood was selected as the best-fitting parameters for each participant.

To evaluate whether both learning rates were necessary, we compared nested models using likelihood ratio tests. The Full model included both **κ**_within_ and **κ**_between_ as free parameters, while three constrained models fixed one or both: (1) a Fixed model with **κ**_within_ = 1 and **κ**_between_ = 0 (no learning from either source), (2) a BlockOnly model with **κ**_within_ = 1 and **κ**_between_ free (learning only from reference presentations), and (3) a TrialOnly model with **κ**_between_ = 0 and **κ**_within_ free (learning only from trial-to-trial experience). For each participant, we extracted the best-fitting parameters for each model from that participant’s own likelihood surface and fitted the corresponding logistic regression. Because the models are nested, we compared them using the likelihood ratio statistic, χ² = −2(LL_constrained - LL_Full), evaluated against a chi-square distribution with degrees of freedom equal to the difference in the number of free parameters between the models (2 for Full vs. Fixed, 1 for the remaining comparisons). This was performed at the individual participant level to assess how many participants required each parameter. At the group level, we computed the sum of the likelihood ratio statistics across all participants, which, under the assumption of independence among observers, follows a chi-square distribution with degrees of freedom equal to the per-subject degrees of freedom multiplied by the number of participants (Claessens & Wagemans, 2008).

### EEG acquisition and preprocessing

EEG was recorded continuously from 64 ActiCap Electrodes at 1000 Hz by a QuickAmp amplifier (Brain Products). Data were high-pass filtered online (0.01 Hz) to avoid electrodermal fluctuations. All electrode sites were referenced to FCz and grounded to AFz. The electrodes were positioned according to the International 10-10 system. Offline, data were re-referenced to the average of TP9 and TP10 (placed on the earlobes). Additional bipolar electrodes recorded the electrooculogram. Data preprocessing and analysis were performed using the Python MNE library. Data was downsampled to 500Hz, and offline filters were applied to the continuous data, using a highpass filter of 0.05 (Butterworth filter, order 2) and a lowpass filter of 30Hz (Butterworth filter, order 4). Channels with missing data due to problems in acquisition or channels with excessive noise were interpolated (mean number of interpolated channels=3, standard deviation = 3). Independent component analysis (ICA) was performed to detect and reject eye-movement artifacts. The ICA was calculated on the continuous data (without epoching) filtered at 1 Hz (Butterworth filter, order 4) using the “fastica” method as implemented in MNE. Eye-related components were identified by visually inspecting topographies and time series from each component. Epochs exceeding ±150 µV at any EEG channel were automatically rejected. Data were epoched relative to interval onset, with a baseline correction applied from −100 to 0 ms. For the analyses reported here, epochs were subsequently realigned to interval offset without recomputing the baseline, preserving the original onset-based baseline correction throughout.

### Multivariate pattern analysis

We investigated how time and category are encoded in time-resolved EEG data using a supervised-learning classification approach. At each time point, all 62 EEG sensors were used as features. We employed Linear Discriminant Analysis (LDA) as implemented in the scikit-learn library (Pedregosa et al., 2011), using a least-squares solution and automatic shrinkage via the Ledoit-Wolf lemma, along with custom scripts. Similar results were obtained using other methods, such as regularized logistic regressions. We restricted our analysis to the three durations (0.5 s, 0.71 s, and 1 s) that, across different blocks, could assume the three possible temporal categories (shorter, equal, and longer). Within these durations, time and category are orthogonal, enabling better estimation of the role of each modulating factor in EEG activity.

Given that both time and category have three possible values, one approach to perform MVPA is to use a multiclass classification. However, this approach would not account for the fact that these values are naturally ordered. For this reason, we employed a binary classification approach, training the classification algorithm on only the two extreme values (0.5 seconds and 1 second for time, and shorter and longer for the category). Critically, the fitted model was subsequently tested on all three values, and the estimated probability for the positive class (1s for time and longer for category) was stored. This score ranges from 0 (low probability of belonging to the positive class) to 1 (high probability of belonging to the positive class). For example, a value of 0.7 indicates that the model predicts that the particular data point belongs to the positive class with 70% probability.

We used a 5-fold cross-validation. Given that our experiment consisted of 100 blocks (20 for each reference), each fold consisted of 4 blocks of each reference. For example, in the first fold, the test data comprised the first four blocks for each reference, and the training data comprised the remaining blocks. Within each cross-validation fold, EEG features were z-scored using the training-set mean and standard deviation, which were then applied to the test set. The analysis was performed at each time point, with samples taken every 2 ms after downsampling. Prior to this decoding analysis, a 50-ms moving-average filter was applied to the epoched data.

### EEG Regression with MVPA

In the second analysis step, we conducted linear regression using the class probability estimates derived from the MVPA. For each participant and time point, the probability of the positive class from the Linear Discriminant Analysis (LDA) classifier served as the dependent variable. These values were regressed on the predictors of interest (e.g., time, category, response), depending on the specific analysis. To account for the bounded nature of probability estimates (0–1), a logit transformation was applied after constraining extreme values to [0.001, 0.999], mapping them onto an unbounded scale. All predictors were standardized (z-scored) before performing ordinary least squares regression. The resulting regression coefficients were stored for subsequent analysis. At the group level, estimated coefficients were compared to zero using a one-sample, two-tailed t-test. Positive and negative values indicate that the particular predictor increased or decreased the estimated probability of the positive class.

We evaluated whether coefficients exceeded chance levels using a permutation-based control of the family-wise error rate for multiple comparisons (Groppe et al., 2011). This method provides strong control of the family-wise error rate (the same degree of false discovery control as Bonferroni’s correction) while generally being more powerful. Permutation-based strong control of the family-wise error rate is one of the most effective methods for establishing reliable lower bounds on the onset and offset of effects when a priori boundaries are unavailable (Groppe et al., 2011). In short, permutations are constructed by flipping the sign of a random set of participants and calculating new *t-*scores under this rearrangement. For each permutation (*n* = 5,000), a *t*_max_ score (the most extreme positive or negative value of the *t* scores across all time points) is stored. Critical *t-*values and *p-*values are estimated from these *t-*max distributions. Only clusters longer than 50ms were considered.

### Temporal generalization

We investigated the cross-temporal dynamics of how category and duration information influenced neural activity. For each participant, we trained LDA classifiers at each time point and tested them at all other time points, creating two-dimensional cross-temporal decoding matrices. We analyzed how predictors (category and target duration) influenced the decoder probability outputs across different training and testing time combinations. For each train-test time pair, decoder probabilities were logit-transformed and regressed on the behavioral predictor of interest (z-scored), as described in the previous section. This yielded a cross-temporal matrix of regression coefficients for each subject. At the group level, we tested whether these coefficients differed significantly from zero using permutation tests as described above. Only cluster sizes of 400 time-point pairs (representing roughly 40 ms × 40 ms of train-test time combinations) were considered.

### Weight Projection

To examine how different sensors contributed to classifier performance, we transformed the classifier weights into activation patterns. This approach provides a more interpretable representation, in which nonzero values indicate class-specific information that can be projected into sensor space. The transformation followed a procedure outlined in a tutorial paper (Grootswagers et al., 2017). Briefly, classifier weights (w) were converted into activation patterns (A) by multiplying them with the covariance matrix of the data: A = cov(X) * w, where X is the N × M matrix of EEG data (with N trials and M features, i.e., channels), and w is a classifier weight vector of length M. The resulting vector A (also of length M) captures the spatial distribution of the informative signal across sensors. For this analysis, classifiers were trained once per timepoint using all available data (without cross-validation) to obtain stable weight estimates. This transformation was performed separately for each participant and time of interest. For visualization, the resulting activation patterns were projected onto the scalp according to the channel locations and then averaged across participants to produce group-level maps.

### Time-Category Pattern Overlap Analysis

To examine potential differences and similarities in how category and duration modulate brain activity, we compared the *weight projections* (estimated as described above) for each dimension (time and category). Weight Projections were estimated in three windows of interest (pre-offset: −200 to −50ms, early: 150 to 300ms, late: 300 to 450ms), and, for each participant and time of interest, we computed the correlation between the category- and duration-classifier weight projections across all 62 EEG channels. High positive correlations indicate that category and duration information are encoded by similar spatial patterns of brain activity, suggesting shared neural substrates. Conversely, correlations near zero indicate that the two dimensions rely on distinct, non-overlapping spatial patterns of neural activity. Negative correlations indicate that regions that contribute positively to one dimension contribute negatively to the other, indicating opposing spatial patterns. Thus, correlations near zero indicates that the two classifiers rely on linearly non-overlapping spatial read-outs, but does not entail independence of the underlying neural generators, which may in principle share common sources while projecting onto distinct axes

### Cross-dimensional generalization

To test whether categorical and temporal information are encoded separately we implemented a leave-one-condition-out cross-validation approach in both directions. First, we trained categorical classifiers on a restricted dataset by excluding all trials from one duration level during training, then tested them on the excluded duration level. For example, we trained categorical classifiers using only trials of 0.5 s and 1.0 s duration, and then tested them on 0.71 s trials. This was repeated for each duration level (0.5 s, 0.71 s, and 1.0 s). Second, we applied the same procedure to duration classifiers: we trained a duration classifier on all categorical contexts except one (e.g., excluding all “longer” trials), then tested on the excluded category. This procedure ensured that every trial served as a test case exactly once, with fold assignments maintained throughout to preserve independence. High correlations between the outputs of the restricted and standard analyses would indicate that classifiers generalize across the excluded dimension, suggesting that the code is preserved across levels of the other variable. Additionally, we examined whether the observed regression effects (e.g., category modulating categorical evidence, duration modulating duration evidence) were preserved in these restricted analyses. If categorical and duration representations are orthogonal, both classifier performance and regression effects should remain stable across excluded levels.

### Statistical Analysis of Decoder Evidence

To examine the factors influencing temporal and decisional signals at specific time points, we performed regression analyses using decoder evidence from the three time windows of interest (pre-offset: −200 to −50ms, early: 150 to 300ms, late: 300 to 450ms) as dependent variables. For each participant, decoder evidence values were logit-transformed after clipping extreme values (0.001-0.999), and linear regression models were fitted with behavioral and experimental predictors (previous response, current category, previous category, target duration, and actual behavioral response) as z-scored independent variables. Group-level inference was performed using one-sample t-tests on regression coefficients across subjects. Given the large number of predictors (15, five per time point of interest), we adjusted for multiple comparisons using the Holm method, applied separately to the categorical and temporal decoding analyses. The scaled Jeffrey-Zellner-Siow (JZS) Bayes Factor was estimated using the implementation in pingouin v.0.5.4 (Vallat, 2018).

## Results

### Behavior

As shown in Figure 1 (panels c-e), participants performed the task well. As expected, the probability of participants responding “longer than reference” was strongly modulated by whether the target was shorter/equal/longer than the reference (1.891 ± 0.097, t(29) = 19.524, p < 0.001, Cohen’s d = 3.565, BF₁₀>1000). Participants’ responses were also modulated by the actual duration of the target, with a higher proportion of longer responses for longer durations (0.355 ± 0.062, t(29) = 5.744, p < 0.001, Cohen’s d = 1.049, BF₁₀ > 1000). Lastly, both the previously presented category and the previous response modulated participants’ current responses, although in opposite directions. The previous category had a negative effect (−0.686 ± 0.045, t(29) = −15.403, p < 0.001, Cohen’s d = −2.812, BF₁₀ > 1000), indicating that the longer the previous category was, the greater the probability that participants responded “short” in the current trial. The previous response, on the other hand, had a positive effect (0.634 ± 0.055, t(29) = 11.546, < 0.001, Cohen’s d = 2.108, BF₁₀ >1000), indicating a tendency to repeat the previous response.

The negative effect of the previous category and the positive effect of target duration suggest that participants might be dynamically updating their temporal reference with both within-block and general across-block information, calibrating their judgments accordingly. To quantify these updating mechanisms, we fitted a model based on the Internal Reference Model to each participant’s data. This model posits that participants maintain an internal reference updated through two mechanisms: (1) within-block updating based on presented trial intervals (**κ**_within_), and (2) between-block updating based on new reference presentations (**κ**_between_). Both parameters range from 0 to 1, with higher values indicating greater retention of the current reference. A theoretically perfect observer would have **κ**_within_ = 1 (perfect retention within blocks) and **κ**_between_ = 0 (complete updating to new references).

Figure 2A shows the distribution of estimated dual-update parameters across participants. The within-block updating rate (**κ**_within_) was significantly higher than the between-block updating rate (**κ**_between_) (**κ**_within: 0.844 ± 0.016, **κ**_between: 0.479 ± 0.033; t(29) = 10.241, p < 0.001, Cohen’s d = 1.87, BF₁₀ > 1000). This pattern suggests a dissociation between the two learning mechanisms: participants maintained good retention of the learned reference within blocks (**κ**_within_ = 0.844 ± 0.016), with modest interference from individual trial intervals, while being more flexible in updating their reference when encountering new reference presentations at the start of each block (**κ**_between_ = 0.479 ± 0.033). These two parameters were uncorrelated (r = 0.072, p = 0.706, BF₁₀ = 0.243), indicating that they capture independent aspects of reference updating.

**Figure 2.**
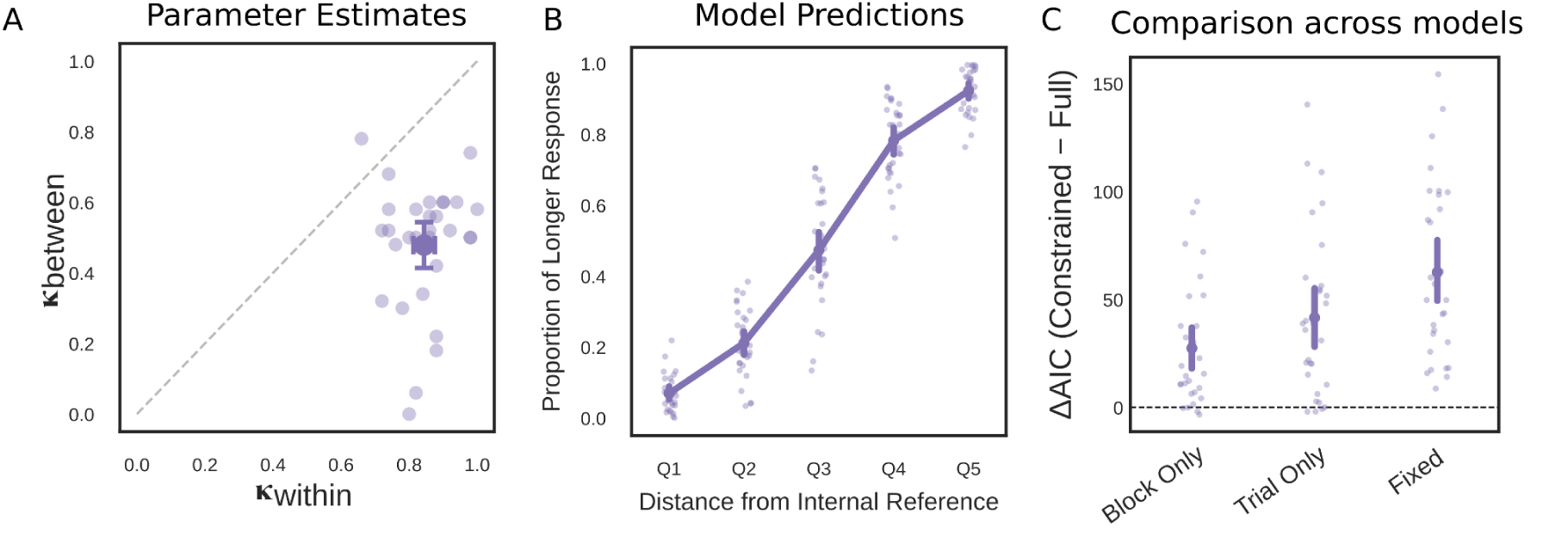
Internal Reference Model results. (A) Distribution of **κ**_within and_ **κ**_between_ across participants. Both parameters range from 0 to 1, with higher values indicating greater retention of the current reference. Individual participants are shown as light dots; the group means are shown with horizontal and vertical error bars representing 95% confidence intervals. (B) Behavioral predictions from the fitted Full model. Proportion of “Longer” responses increases as a function of distance from the continuously updated internal reference (quintiles Q1–Q5). Individual participant data are shown as light dots with substantial variability, while the solid line with error bars represents group means and 95% confidence intervals. (C) Model comparison across constrained alternatives. Each point represents one participant’s ΔAIC (Constrained - Full) for the BlockOnly, TrialOnly, and Fixed models. Positive values indicate better fit for the Full model after penalizing for additional parameters. Group means are shown with 95% confidence intervals

If participants dynamically update their internal reference both within and across blocks, the Full model incorporating both κ_within and κ_between should outperform models with either parameter fixed. The likelihood ratio tests indicated that both learning rates were necessary to account for participants’ behavior. At the individual level, the Full model significantly outperformed the Fixed model (χ²(2), p < .05) across all 30 participants (median χ²= 57.46), indicating that each participant benefited from at least one form of reference updating. The Full model also significantly outperformed the BlockOnly model (χ²(1), p < .05) in 25 of 30 participants (median χ²= 19.37). Similarly, the Full model outperformed the TrialOnly model (χ²(1), p < .05) in 26 of 30 participants (median χ² = 39.32). Group-level summed likelihood ratio tests corroborated these findings, with the Full model significantly outperforming all constrained models (Full vs. Fixed: χ²(60) = 1997.71, p < .001; Full vs. BlockOnly: χ²(30) = 882.45, p < .001; Full vs. TrialOnly: χ²(30) = 1306.84, p < .001). These results suggest that observers employ a dual-timescale updating strategy. Within blocks, participants maintain stable reference representations while resisting interference from individual comparison intervals, and across block boundaries, they update toward new references while retaining some influence from prior experience rather than fully resetting.

### Multivariate pattern analysis

If duration and categorical decision information are encoded by distinct neural processes, we expect to find: (1) different temporal profiles for the two signals; (2) spatially non-overlapping activation patterns; and (3) classifiers that generalize across levels of the orthogonal variable. To test for these possibilities, we applied multivariate pattern analysis (MVPA) to the EEG data surrounding interval offset. Unlike previous studies, we did not apply baseline correction to the EEG signal, allowing us to examine both pre- and post-offset activity without distortion. We trained separate Linear Discriminant Analysis (LDA) classifiers to decode duration information and categorical decisions at each time point. Classifiers were trained on all trials (correct or incorrect) but only at the extreme levels of each feature (0.5s vs. 1s for duration and shorter vs. longer for categorical decisions) to maximize discriminability while avoiding assumptions about intermediate categories. However, they were applied to all levels at the testing phase. The classifier outputs—probability estimates for the positive class—were then subjected to regression analysis to examine how experimental factors modulated neural representations over time. Unlike conventional MVPA approaches that report classifier performance as accuracy or AUC, here classifier performance is evaluated by regressing the logit-transformed probability estimates from the LDA classifier against the experimental variables of interest, and reporting the resulting beta coefficients. This approach allows us to assess not only whether a variable is decodable, but also how strongly and in which direction it modulates the classifier output, and to investigate, in subsequent analyses, what experimental and behavioral factors modulate the decoded signal.

Decoder performance was evaluated by computing the decoder probability across all test trials and regressing it against the true class. As shown in Figure 3, decoding of target duration and categorical variables showed different time courses. Although both types of information could be decoded during the entire offset and post-offset period, duration information was more strongly encoded during the pre-offset period, while categorical decisions were more prominent after the interval offset. To examine the temporal dynamics and stability of duration and categorical representations, we performed a temporal generalization analysis (Figure 3). Classifiers were trained at each time point and tested on all other time points, generating cross-temporal decoding matrices that reveal how neural representations evolve and generalize over time. The temporal generalization patterns revealed a similar time-specific pattern of generalization, with stronger performance for duration in the pre-offset period and for category post offset, suggesting that both representations are dynamically evolving and time-locked to specific phases of interval processing.

**Figure 3.**
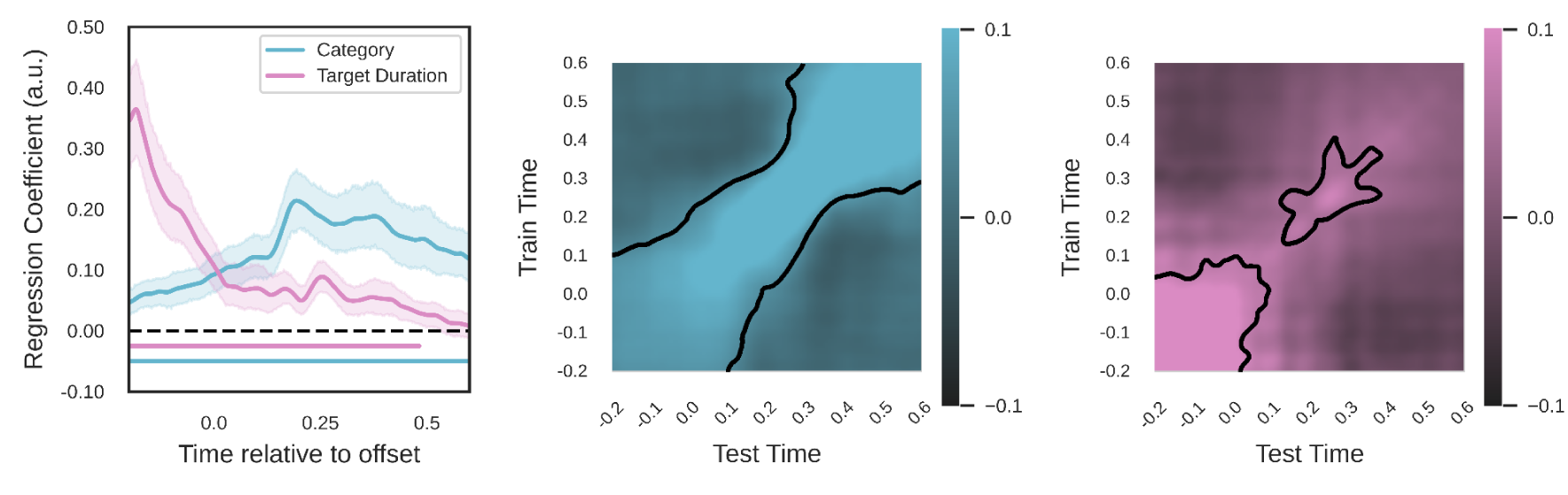
Time-resolved decoding and temporal generalization of duration and category information. (Left) Decoding of category (blue) and target duration (pink). Duration information is more strongly encoded during the pre-offset period, while categorical decisions become more prominent after the interval offset. The horizontal lines below indicate periods of statistical significance (permutation test, p < 0.05). (Center and Right) Cross-temporal generalization matrices showing classifier performance when trained at one time point (y-axis) and tested at another (x-axis). (Center) Category information exhibits strong decoding performance, particularly along the diagonal, suggesting dynamically evolving representation that are time-locked to specific processing phases. (Right) Duration information exhibits more transient, time-specific patterns with stronger concentration during the pre-offset period. Significant clusters are identified using permutation testing under the null hypothesis of zero.

Based on these MVPA results, we identified three moments of interest for subsequent analyses: (1) a pre-offset window where duration information was strongly represented, (2) an early post-offset window showing robust categorical encoding, and (3) a late post-offset window where both signals remained detectable. These windows were selected to capture the distinct phases of temporal information processing revealed by the multivariate decoding analyses.

### Time-Category Pattern Overlap

Weight pattern correlations provided evidence for spatial independence. Direct correlations between the topographic patterns of category and duration classifiers were consistently near zero across time windows (Pre-offset: r = 0.099 ± 0.082, t = 1.205, p = 0.238, d=0.220; Early: r = −0.058 ± 0.073, t = −0.795, p = 0.433, d = 0.145; Late: r = 0.087 ± 0.073, t = 1.196, p = 0.242, d=0.218), with all correlations falling within narrow confidence intervals that include zero (95% CIs: [-0.062, 0.259], [-0.201, 0.085], and [-0.056, 0.230] respectively). Bayesian analyses provided moderate-to-strong evidence for the null hypothesis of zero correlation (BF₁₀ = 0.376, 0.260, and 0.372), supporting the spatial independence of duration and categorical encoding patterns.

To test further whether time and category are encoded separately, we employed a leave-one-condition-out approach, training classifiers on data excluding one level of the control variable (e.g., training category classifiers without trials of 0.71 seconds and testing on the excluded level). The resulting classifier outputs showed strong correlations with the standard analysis that included all conditions, indicating that the learned decision boundaries generalize across different levels of the orthogonal dimension. For category decoding, correlations were high in each time window (Pre-offset: r = 0.795 ± 0.009; Early: r = 0.846 ± 0.007; Late: r = 0.841 ± 0.008; all p < 0.001). Duration decoding showed similarly strong correlations (Pre-offset: r = 0.874 ± 0.006; Early: r = 0.826 ± 0.004; Late: r = 0.822 ± 0.005; all p < 10⁻⁴²). These high correlations (all r > 0.79), large effect sizes (Cohen’s d > 16), and Bayesian evidence (BF₁₀ > 1000) are consistent with categorical and temporal representations generalizing across different levels of the orthogonal dimension. These high correlations, however, should be interpreted with caution, given that roughly two-thirds of the training data is shared between the standard and restricted classifiers, which naturally increases their magnitude. On the other hand, the excluded third is not randomly chosen, as it consists of all trials from one level of the orthogonal dimension. A classifier that relied heavily on the orthogonal variable to discriminate categories would fail to generalize well to these excluded trials.

To validate the separability between duration and categorical representations, we conducted regression analyses to examine how experimental variables modulate classifier evidence. For each participant and time window, we regressed the logit-transformed classifier probabilities against the relevant predictors (category for category classifiers, target duration for duration classifiers). We compared regression coefficients from the standard analysis (trained on all conditions) with those from the orthogonality test (trained excluding one level of the control variable) to test whether the experimental effects remain consistent when classifiers are trained on reduced datasets that exclude specific levels of the orthogonal dimension. Both standard and orthogonal analyses showed significant modulation by the relevant experimental variables, with effects preserved across training conditions.

For category classifiers, category significantly modulated neural evidence across all time windows in both standard (Pre-offset: β = 0.076 ± 0.014, t(29)=5.314, p<0.001, d=0.970, BF>1000; Early: β = 0.190 ± 0.023, t(29)=8.427, p<0.001, d=1.539, BF>1000; Late: β = 0.185 ± 0.027, t(29)=6.826, p<0.001, d=1.246, BF>1000) and orthogonal analyses (Pre-offset: β = 0.059 ± 0.017, t(29)=3.491, p=0.002, d=0.637, BF=22.4; Early: β = 0.175 ± 0.025, t(29)=7.033, p<0.001, d=1.284, BF>1000; Late: β = 0.172 ± 0.030, t(29)=5.803, p<0.001, d=1.059, BF>1000). The coefficients showed the strongest effects in the Early and Late time windows, consistent with post-offset categorical processing.

For duration classifiers, target duration significantly modulated evidence in both standard (Pre-offset: β = 0.282 ± 0.033, t(29)=8.553, p<0.001, d=1.562, BF>1000; Early: β = 0.068 ± 0.012, t(29)=5.415, p<0.001, d=0.989, BF>1000; Late: β = 0.056 ± 0.013, t(29)=4.195, p<0.001, d=0.766, BF>1000) and orthogonal conditions (Pre-offset: β = 0.268 ± 0.034, t(29)=7.969, p<0.001, d=1.455, BF>1000; Early: β = 0.055 ± 0.013, t(29)=4.208, p<0.001, d=0.768, BF>1000; Late: β = 0.044 ± 0.014, t(29)=3.179, p=0.004, d=0.580, BF=11.1). Duration effects were strongest during the Pre-offset period. The regression coefficients remained highly similar between the standard and orthogonal analyses, with strong Bayesian evidence supporting the presence of experimental effects in both conditions.

To examine the spatial organization of duration and categorical representations, we plotted mean classifier outputs for each combination of target duration (0.5s, 0.71s, 1.0s) and category (shorter, equal, longer) in a two-dimensional space, with category evidence on the x-axis and duration evidence on the y-axis (Figure 4). Data points formed a systematic grid pattern across all time windows, with category information varying horizontally (represented by different colors) and duration information varying vertically (represented by different markers). The orthogonal arrangement of conditions in this neural decision space suggests that duration and categorical information are encoded along spatially separable dimensions. Points representing different durations were separated along the y-axis regardless of their categorical status, while points representing different categories were separated along the x-axis regardless of their physical duration. This orthogonal organization was consistent across time windows, indicating that the brain simultaneously encodes both physical duration and categorical temporal decisions along separate neural dimensions.

**Figure 4.**
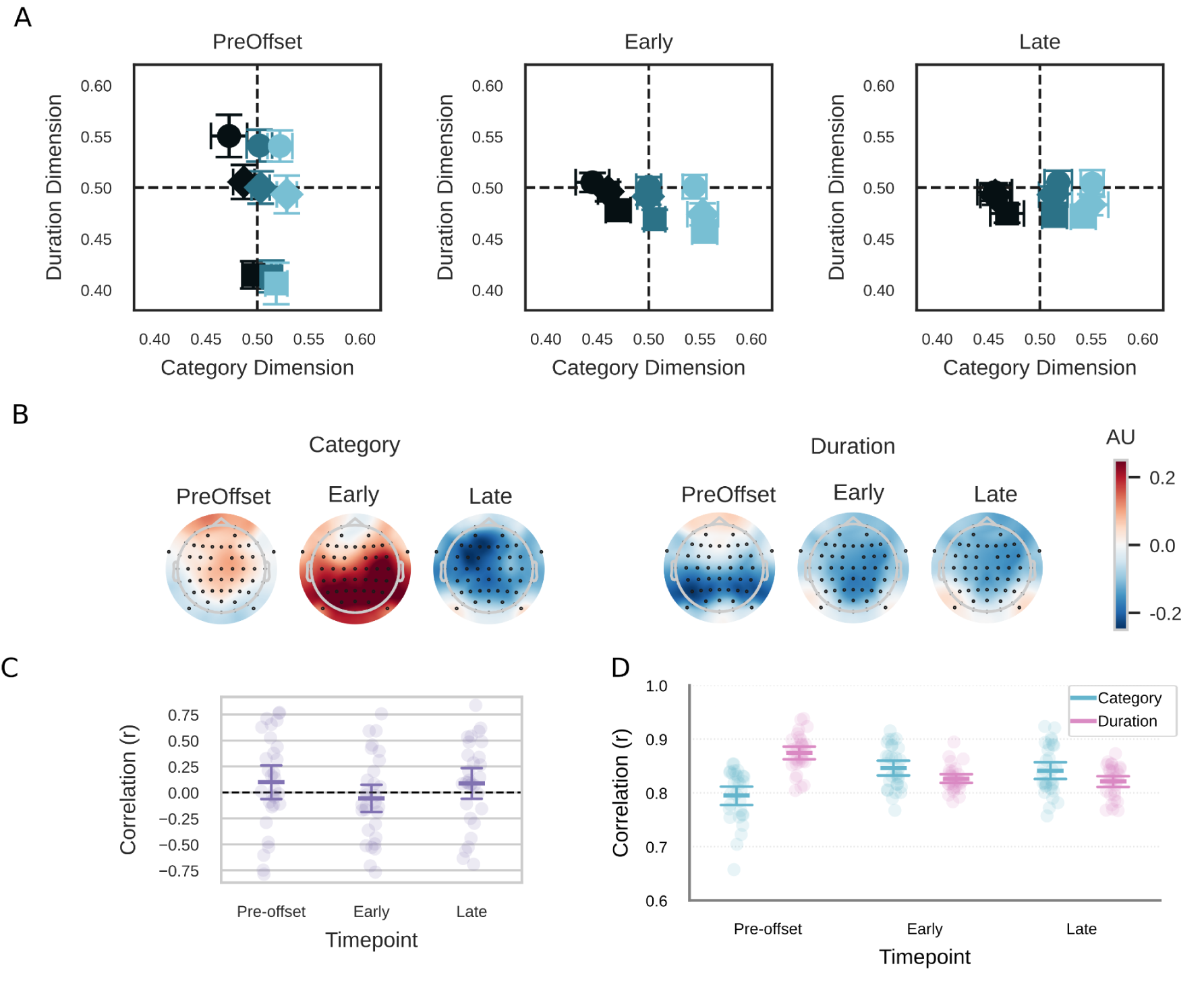
Separable encoding of duration and category information in neural decision space. (A) Two-dimensional visualization of neural representations across time windows, with category classifier evidence on the x-axis and duration classifier evidence on the y-axis. Each point represents the mean classifier output for a specific combination of target duration (markers: square = 0.5s, diamond = 0.71s, circle = 1.0s) and category (colors: dark = shorter, medium = equal, light = longer than reference), with error bars showing 95% confidence intervals. Reference lines at 0.5 indicate chance-level classification. The systematic grid-like organization in panel A provides an intuitive illustration of separable encoding: if duration and category were encoded along shared dimensions, points representing different durations within the same category (or different categories within the same duration) would overlap rather than forming distinct rows and columns. (B) Spatial activation patterns (weight projections) for category and duration classifiers across time windows. Topographic maps show the scalp distribution of discriminative information, with red indicating positive weights and blue indicating negative weights. (C) Orthogonality analysis showing correlations between weight patterns (right). Correlations remain near zero. Individual participant data are shown as points, group means with 95% confidence intervals. (D) Cross-validation orthogonality test showing correlations between standard classifier outputs (trained on all conditions) and orthogonal classifier outputs (trained excluding one level of the control variable). Individual participant data are shown as points, group means with 95% confidence intervals.

### Experimental and Behavioral Predictors of Classifier Evidence

To investigate how experimental and behavioral factors affect neural representations of duration and category information, we conducted regression analyses using these features as predictors. For each participant and time window (pre-offset, early, and late), we regressed logit-transformed classifier evidence against standardized predictors, including category, target duration, previous category, previous response, and current response. The regression analysis revealed systematic influences of experimental variables on neural representations, with different predictors showing varying effects across time windows and between category and duration classifiers (Figure 5).

**Figure 5.**
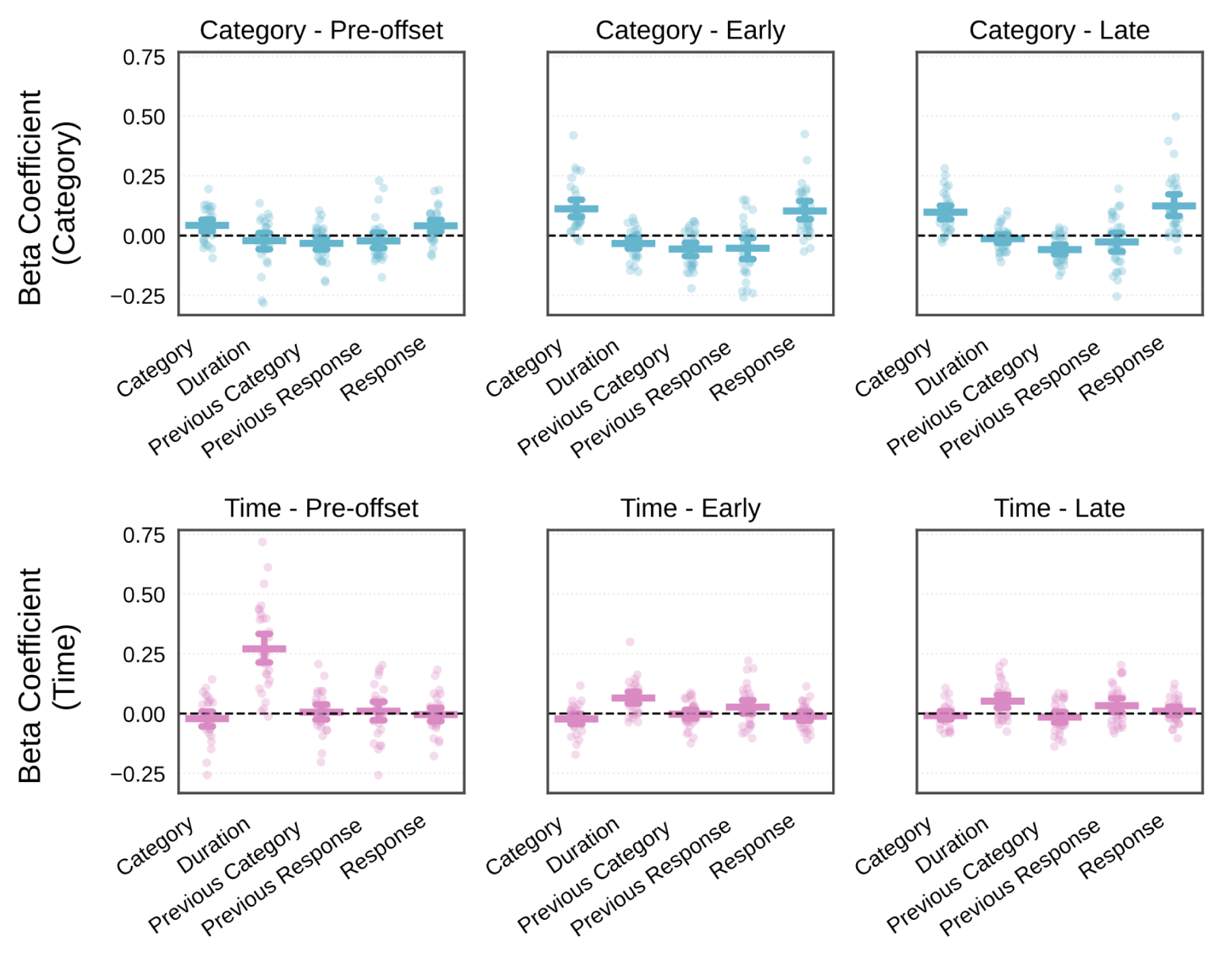
Regression analysis of experimental and behavioral predictors on classifier evidence. Beta coefficients from linear regressions examining how multiple factors modulate neural evidence for category (top row) and duration (bottom row) classifiers across three time windows relative to interval offset. Each subplot shows individual participant data (points) and group means with 95% confidence intervals (horizontal lines). Predictors include categorical status (Category), target duration (Duration), previous trial category (Previous Category), previous motor response (Previous Response), and current behavioral response (Response). Category classifiers exhibit the strongest modulation by categorical variables and behavioral responses, with a temporal evolution from weak pre-offset to strong post-offset effects. Duration classifiers show selective and consistent modulation by target duration across all time windows, with minimal influence from other variables. Dashed horizontal lines indicate zero (no effect).

The category classifier had a complex pattern of modulations by different predictors. As expected, the objective category (shorter/equal/longer than reference) showed the strongest and most consistent effects across all time windows, with significant modulation increasing from pre-offset to post-offset periods (Pre-offset: β = 0.043 ± 0.012, p = 0.017, d = 0.625, BF_10_ = 19.11; Early: β = 0.112 ± 0.019, p < 0.001, d = 1.093, BF_10_ > 1000; Late: β = 0.098 ± 0.015, p < 0.001, d = 1.182, BF_10_ > 1000). This pattern reflects the temporal dynamics of categorical decision formation, with weak early encoding that strengthens after the interval offset. Previous category did not modulate category evidence during the pre-offset period (β = −0.032 ± 0.013, p = 0.112, d = −0.450, BF_10_ = 2.53) but significantly influenced post-offset processing (Early: β = −0.056 ± 0.014, p = 0.004, d = −0.726, BF10 = 70.62; Late: β = −0.059 ± 0.010, p < 0.001, d = −1.035, BF_10_ > 1000). The negative coefficients indicate contrast effects consistent with dynamic reference updating that emerges after stimulus presentation. Previous response did not significantly influence category evidence at any time window (Pre-offset: p = 0.738, d = −0.248, BF_10_ = 0.45; Early: p = 0.112, d = −0.455, BF_10_ = 2.67; Late: p = 0.738, d = −0.245, BF_10_ = 0.44), indicating that motor history had a minimal impact on categorical representations.

Behavioral response prediction revealed a strong coupling between neural evidence and decision output. The actual behavioral response (whether participants judged that particular interval as longer or shorter) significantly modulated category classifier evidence across all time windows (Pre-offset: β = 0.041 ± 0.012, p = 0.021, d = 0.602, BF_10_ = 14.39; Early: β = 0.103 ± 0.019, p < 0.001, d = 0.966, BF_10_ > 1000; Late: β = 0.124 ± 0.023, p < 0.001, d = 0.972, BF_10_ > 1000), with effects strengthening over time.

Target duration did not modulate category evidence during pre-offset or late periods (Pre-offset: p = 0.738, d = −0.224, BF_10_ = 0.39; Late: p = 0.738, d = −0.245, BF_10_ = 0.44), but showed significant influence during the Early time window (β = −0.033 ± 0.011, p = 0.041, d = −0.543, BF_10_ = 7.11). This is consistent with the behavioral finding that target duration can also modulate participants’ responses even after accounting for categorical status, as captured by the Internal Reference Model.

Duration classifier evidence was primarily and consistently modulated by target duration across all time windows. Actual target duration showed the strongest effects on duration classifiers, with significant modulation that was strongest during the pre-offset period and remained throughout (Pre-offset: β = 0.271 ± 0.033, t = 8.160, p < 0.001, d = 1.490, BF_10_ > 1000; Early: β = 0.065 ± 0.013, t = 5.030, p < 0.001, d = 0.918, BF_10_ = 972.78; Late: β = 0.051 ± 0.014, t = 3.722, p = 0.011, d = 0.679, BF_10_ = 38.38). This pattern reflects the expected temporal dynamics of duration processing, with the strongest decoding occurring during stimulus accumulation and persisting into post-offset periods with diminishing strength.

Cross-dimensional effects from categorical variables were absent across all time periods. Categorical status did not significantly modulate duration classifier evidence at any time window (Pre-offset: β = −0.021 ± 0.016, t = −1.306, p = 1.000, d = −0.238, BF_10_ = 0.42; Early: β = −0.023 ± 0.011, t = −2.067, p = 0.526, d = −0.377, BF_10_ = 1.24; Late: β = −0.009 ± 0.009, t = −1.043, p = 1.000, d = −0.190, BF_10_ = 0.32), demonstrating relative independence of duration processing from categorical decision variables.

Trial history effects were minimal and non-significant for duration processing. Previous category did not influence duration evidence at any time window (Pre-offset: β = 0.005 ± 0.016, t = 0.351, p = 1.000, d = 0.064, BF_10_ = 0.21; Early: β = −0.003 ± 0.010, t = −0.258, p = 1.000, d = −0.047, BF_10_ = 0.20; Late: β = −0.015 ± 0.012, t = −1.302, p = 1.000, d = −0.238, BF_10_ = 0.42), indicating that categorical trial history does not carry over to duration representations. Previous response showed no significant effects (Pre-offset: β = 0.010 ± 0.020, t = 0.505, p = 1.000, d = 0.092, BF_10_ = 0.22; Early: β = 0.027 ± 0.015, t = 1.785, p = 0.847, d = 0.326, BF_10_ = 0.79; Late: β = 0.033 ± 0.015, t = 2.189, p = 0.442, d = 0.400, BF_10_ = 1.53), though there was a trend toward positive influence during later time periods that did not reach significance after correction. Lastly, behavioral response coupling was absent for duration classifiers. The actual behavioral response did not significantly modulate duration classifier evidence at any time window (Pre-offset: β = −0.004 ± 0.014, t = −0.305, p = 1.000, d = −0.056, BF_10_ = 0.20; Early: β = −0.011 ± 0.010, t = −1.171, p = 1.000, d = −0.214, BF_10_ = 0.36; Late: β = 0.010 ± 0.009, t = 1.051, p = 1.000, d = 0.192, BF_10_ = 0.32). This absence of coupling between duration evidence and categorical decisions further supports the independence of duration processing from decision-making variables.

These findings suggest that duration classifiers selectively encode temporal information while remaining largely separable of categorical variables, trial history, and decision outcomes.

### The relation between pattern activity and IRM

If the category classifier tracks a continuously updated decisional signal rather than discrete category membership, its output should be better predicted by the signed distance from the participant’s dynamically updated internal reference than by the nominal category label, and should show systematic modulation even on equal trials where the nominal label carries no information. The modulation of category classifier evidence by previous trial category, reported above, is consistent with this possibility. To test it directly, we conducted a model comparison analysis. For each participant, we fit two competing linear regression models to predict category-classifier evidence (logit-transformed) at each time window (Pre-offset, Early, Late). The first model used the true categorical status (short/same/long, coded as −1/0/1) as the predictor variable, whereas the second model used the signed distance from the internal reference as the predictor variable. Both predictors were z-scored within each participant to ensure comparable scaling. We then compared model fit using the coefficient of determination (R^2^), which quantifies how well each model accounts for the observed data; higher values indicate better fit.

Results showed that the model using distance from the internal reference explained more variance in category-classifier evidence than the simple category label model across all time periods. During the Pre-offset period, the distance predictor showed numerically higher R² compared to the categorical predictor, though this difference was not significant (ΔR² = 0.001 ± 0.000, t(29) = 1.885, p = .070, d = 0.344, BF₁₀ = 0.92), with 17 of 30 participants favoring the distance model. In the Early post-offset period, the distance predictor outperformed the category label (ΔR² = 0.003 ± 0.001, t(29) = 2.204, p = .036, d = 0.402, BF₁₀ = 1.57), with 21 of 30 participants favoring the distance model. This pattern persisted into the Late period (ΔR² = 0.003 ± 0.001, t(29) = 2.310, p = .028, d = 0.422, BF₁₀ = 1.90), with 19 of 30 participants showing higher R² for the distance predictor. Although per-window evidence was anecdotal (BF₁₀ = 0.92–1.90), the distance model was numerically favored in all three windows and by a majority of participants in each.

To provide a more direct test, we restricted the analysis to equal trials, in which the nominal category label has no variance and therefore cannot predict classifier evidence by construction. We computed the within-participant correlation between IRM distance and logit-transformed classifier evidence on these trials, and tested the resulting correlations at the group level. As shown in Figure 6, IRM distance reliably predicted classifier evidence in all three windows, with strong evidence in the post-offset period (Pre-offset: r = 0.027 ± 0.010, t(29) = 2.694, p = .012, d = 0.492, BF₁₀ = 3.97, 22/30 participants with r>0; Early: r = 0.057 ± 0.014, t(29) = 4.220, p < .001, d = 0.771, BF₁₀ = 127.95, 23/30 participants with r>0; Late: r = 0.054 ± 0.015, t(29) = 3.625, p = .001, d = 0.662, BF₁₀ = 30.57, 22/30 participants with r>0). Together, these results suggest that category classifier evidence reflects a graded, continuously updated decisional signal rather than discrete category membership, with the equal-trial analysis providing the clearer and stronger evidence for this interpretation.

**Figure 6.**
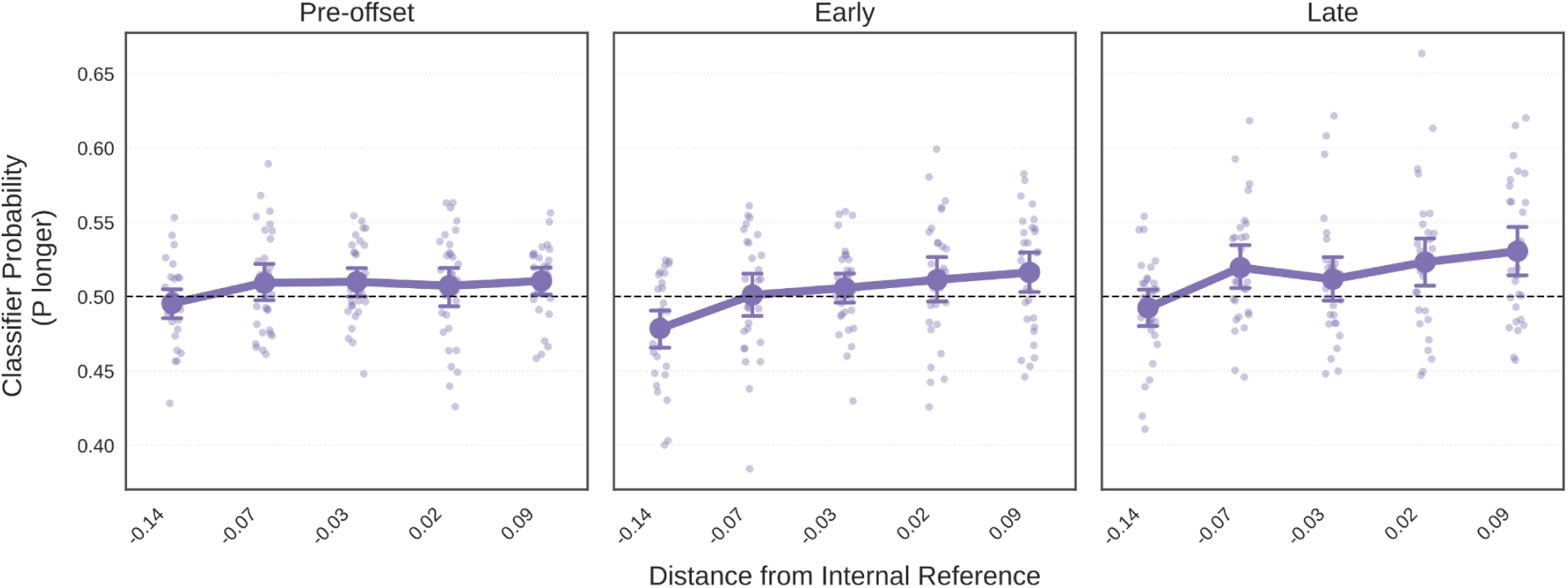
Category classifier evidence on equal trials as a function of distance from internal reference. Classifier probability (P longer) on equal trials (target duration matched the block reference) as a function of binned distance from the participant’s dynamically updated internal reference, shown separately for the three time windows of interest (Pre-offset, Early, Late). Negative values indicate targets shorter than the internal reference; positive values indicate targets longer than the internal reference. Points show individual subject means; lines show group means with 95% CIs (N=30). A monotonic increase from negative to positive IRM distance, most prominent in the Early and Late post-offset windows, indicates that classifier evidence tracks a graded decisional signal even on trials where the nominal category label carries no information.

### Exploratory Event-Related Potentials Analysis

The preceding analyses employed multivariate pattern analysis (MVPA), which offers several methodological advantages: it leverages the multivariate structure of neural signals and enables a more data-driven analysis. However, given recent interest in event-related potentials (ERPs) associated with temporal processing, and to facilitate comparison with prior ERP studies, we conducted supplementary analyses examining univariate ERP responses (ERPs time-locked to the interval onset can be seen in Figure S1).

Because the temporal windows and electrode clusters were selected based on the observed MVPA results, standard inferential statistics would be inappropriate due to circularity in the analysis stream. Therefore, we adopted an exploratory approach to characterize how category, target duration, and distance from the internal reference modulated ERP amplitudes within these predefined spatiotemporal regions of interest.

To characterize the relationship between ERP amplitude and temporal variables, we computed within-subject Pearson correlations for three predictors: signed temporal category (−1/0/+1 for Shorter/Same/Longer), absolute category distance, and target duration. Correlations were computed separately for each of the three MVPA-identified spatiotemporal regions of interest: (1) central electrodes (F1, Fz, F2, FC1, FC2, C1, Cz, C2) during the pre-offset period, (2) parieto-occipital electrodes (PO3, POz, PO4, PO7, PO8, O1, Oz, O2) during the early post-offset period, and (3) central electrodes during the late post-offset period.

For the central pre-offset window (Figure 7), the signed category showed a weak positive correlation with ERP amplitude (r = 0.035 ± 0.012, 95% CI [0.011, 0.060], d = 0.52), indicating more positive ERPs for longer than for shorter categories. Absolute category distance also showed a weak positive correlation (r = 0.022 ± 0.010, 95% CI [0.002, 0.042], d = 0.40), while target duration showed virtually no relationship with amplitude (r = −0.001 ± 0.014, 95% CI [-0.029, 0.027], d = −0.01).

**Figure 7.**
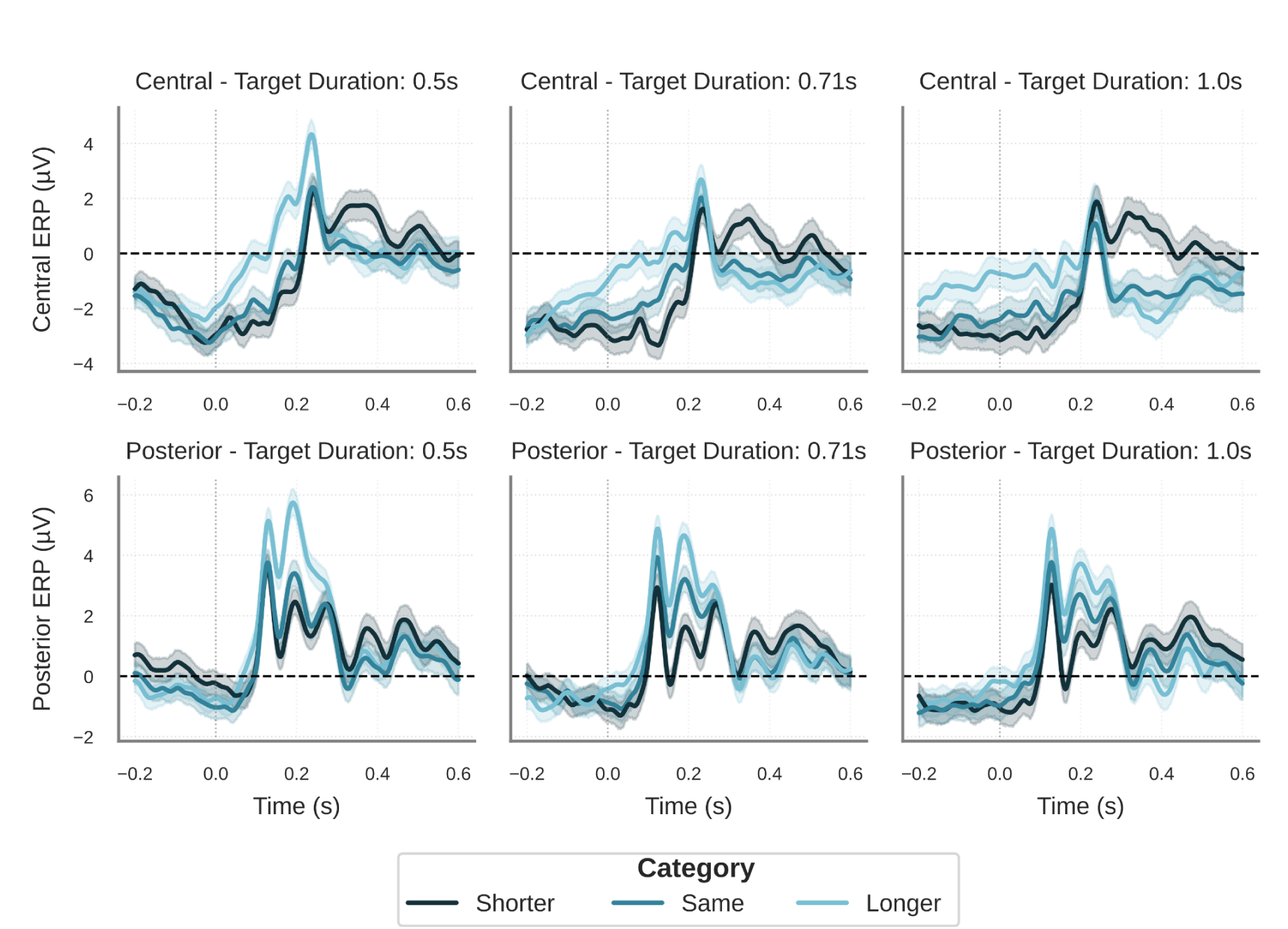
Event-related potentials in central and posterior ROIs as a function of target duration and temporal category. ERPs are shown for central (top row) and posterior (bottom row) regions of interest, separately for each target duration (0.5s, 0.71s, and 1.0s; columns). Data are aligned to stimulus offset (time 0, indicated by the gray vertical dotted line). The three temporal categories (Shorter, Same, Longer) are shown in different shades, representing the target duration relative to the participant’s internal reference. Lines represent the mean across participants, and shaded areas indicate 95% confidence intervals obtained via bootstrap resampling (n=1000).

The posterior early window exhibited the strongest category effect, with signed category showing a moderate positive correlation (r = 0.090 ± 0.014, 95% CI [0.063, 0.117], d = 1.21), suggesting more positive ERPs for longer categories. Absolute category distance showed minimal correlation (r = 0.010 ± 0.010, 95% CI [-0.010, 0.030], d = 0.19), while target duration showed a weak negative correlation (r = −0.028 ± 0.010, 95% CI [-0.047, −0.009], d = −0.53).

In the central late window, signed category showed a negative correlation (r = −0.066 ± 0.018, 95% CI [-0.101, −0.031], d = −0.68), indicating more negative ERPs for longer categories, a reversal of the earlier positive relationship. Absolute category distance showed a weak positive correlation (r = 0.027 ± 0.010, 95% CI [0.007, 0.047], d = 0.48), and target duration showed a weak negative correlation (r = −0.049 ± 0.008, 95% CI [-0.065, −0.033], d = −1.10).”

We next examined how the distance from each participant’s internal reference (i.e., the difference between the target duration and the internal reference) related to ERP amplitude (Figure 8). We computed within-subject Pearson correlations for two measures: signed distance (negative values indicate targets shorter than the internal reference, positive values indicate longer targets) and absolute distance (the magnitude of the deviation regardless of direction).

**Figure 8.**
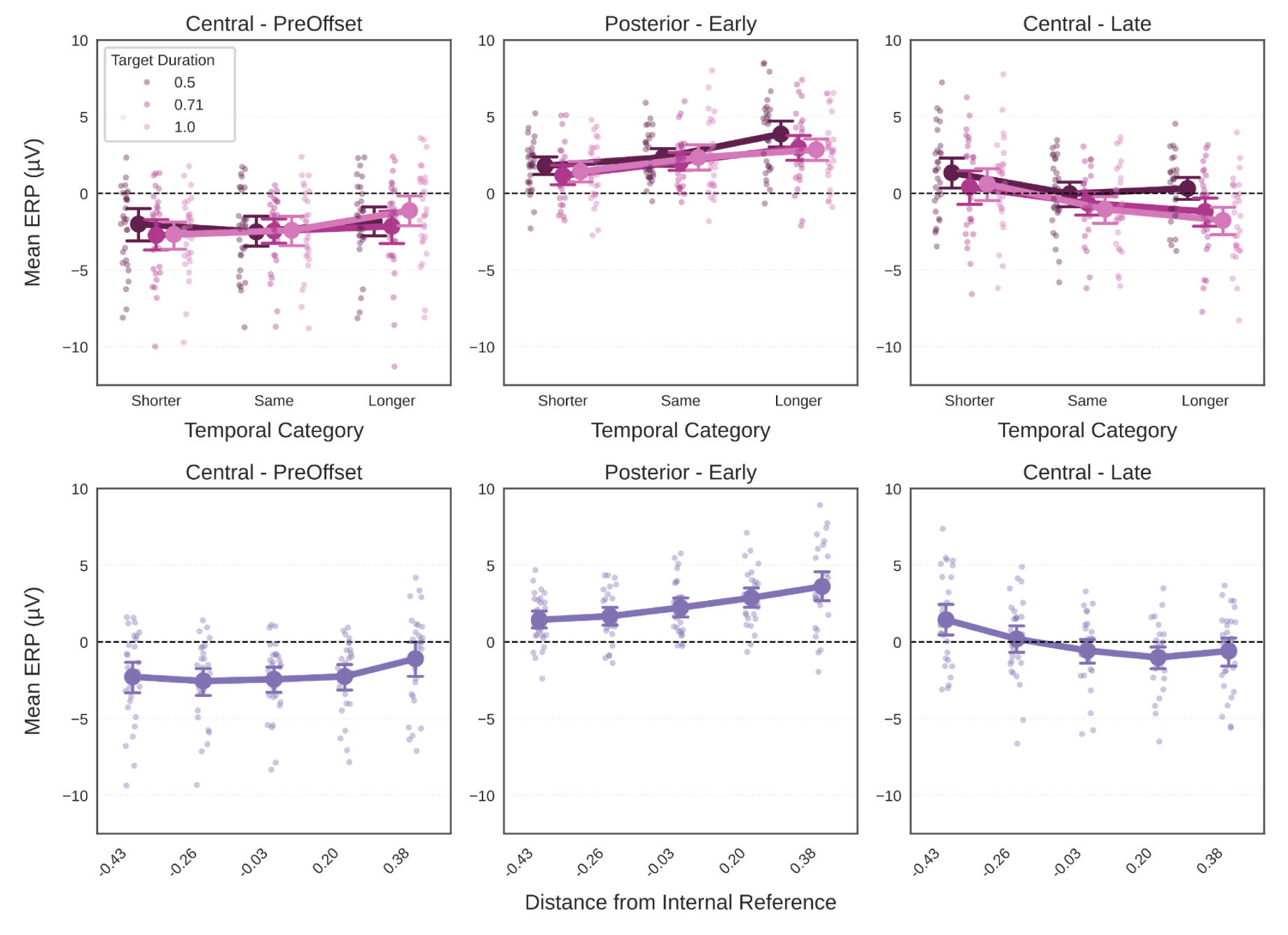
ERP amplitudes by temporal category, target duration, and internal reference distance. Top: ERPs for temporal categories (Shorter, Same, Longer) across three target durations (0.5s, 0.71s, 1.0s) in central pre-offset (left), posterior early (middle), and central late (right) time windows. Bottom: ERPs as a function of distance from internal reference for the same ROIs/time windows. Points show individual subject means; lines show group means with 95% CIs (N=30). Time windows and ROIs were selected based on MVPA results.

For the central pre-offset window, signed distance showed a weak positive correlation with ERP amplitude (r = 0.039 ± 0.013, 95% CI [0.014, 0.065], d = 0.56), indicating more positive ERPs for targets longer than the internal reference. Absolute distance also showed a weak positive correlation (r = 0.029 ± 0.010, 95% CI [0.009, 0.049], d = 0.52), suggesting greater ERP amplitudes for larger deviations from the internal reference.

The parieto-occipital early window showed the strongest signed distance effect, with a moderate positive correlation (r = 0.092 ± 0.015, 95% CI [0.063, 0.121], d = 1.13), indicating more positive ERPs for targets exceeding the internal reference. However, absolute distance showed virtually no correlation (r = −0.002 ± 0.009, 95% CI [-0.020, 0.017], d = −0.03), suggesting that the signed relationship, rather than mere distance magnitude, drove the effect.

In the central late window, signed distance showed a weak negative correlation (r = −0.069 ± 0.018, 95% CI [-0.104, −0.035], d = −0.71), representing a polarity reversal relative to earlier windows. Absolute distance showed a weak positive correlation (r = 0.048 ± 0.011, 95% CI [0.027, 0.069], d = 0.80), indicating that larger deviations from the internal reference, regardless of direction, were associated with more positive ERPs in this late processing stage

## Discussion

We used a design in which physical duration and category (shorter, equal, or longer than a reference) varied independently to investigate their neural representations. We found three main results. First, participants updated their internal temporal reference both trial-by-trial and block-by-block. Second, duration and categorical information showed different profiles: duration peaked before offset, whereas categorical decisions peaked after offset. Third, these signals were represented along separable dimensions.

Our behavioral results show that temporal judgments were strongly shaped by recent experience. Participants’ responses were modulated by current target duration and category, as well as by the category presented on the previous trial and the previous response. To model the updating process, we used a variation of the Internal Reference Model (IRM) with a between-block learning mechanism. The model captured both within-block adaptation and cross-block persistence of learned temporal contexts, indicating that temporal memory can operate on multiple timescales.

In most previous temporal categorization studies, duration and category covary, making it difficult to isolate activity reflecting the encoding of duration and decision variables. Our orthogonal design separated these factors, revealing that they are represented along different activity patterns. This separability was evidenced by the lack of consistent similarities between the topographies of the two effects and by the generalization of decoding across different levels of both variables. Duration information peaked before stimulus offset, whereas categorical decision signals emerged primarily after offset. Temporal generalization indicated that both were transient and time-specific, suggesting a dynamic evolution. A methodological point worth addressing is that our classifiers were trained on the two extreme durations (0.5 and 1s). These durations likely fall on different sides of a perceptual boundary (Grondin, 2012; Grondin et al., 2004; Lewis & Miall, 2003; Thibault et al., 2023, 2024). Our data are consistent with this proposal, as duration was decoded from neural activity. Because these two training durations are likely not perceptually interchangeable, the emergence of a consistent categorical signal separable from the duration signal reinforces the proposal of a temporal decisional signal that is consistent across different durations.

Activity associated with categorical information was observed both before and after interval offset, indicating the presence of categorical judgments during ongoing temporal accumulation. This pre-offset categorical signal generalized across different physical durations, suggesting that neural activity was encoding the target’s categorical relationship to the reference rather than its duration. This finding speaks to a longstanding debate about whether pre-offset activity, typically measured as the contingent negative variation (CNV), reflects purely temporal accumulation or already incorporates decision-related information. Our results suggest that categorical decision signals are present during the pre-offset period and are separable from duration-related activity. Instead of relying on a single ERP component measured at pre-selected electrodes, we leveraged the full multivariate structure of the EEG to identify broad-scale activity patterns that tracked the target category. This approach is more sensitive to distributed neural representations and free from assumptions about where in the scalp the relevant signal should arise. One caveat concerns the pre-offset duration signal specifically: because the pre-offset window is defined relative to offset, time-since-onset is perfectly confounded with physical duration. Crucially, our orthogonal design breaks the confound between elapsed time and decision, since the same duration maps onto different categorical positions across blocks.

During the post-offset period, categorical information became dominant, whereas duration information declined, suggesting a shift toward explicit categorization. This increase may reflect several factors. First, the stimulus offset itself serves as a discrete event that triggers a cascade of decision-related processes, memory comparison, category assignment, and response selection, each contributing additional variance to the signal (Bueno et al., 2024). Second, with the temporal information now complete, post-offset activity may reflect a more resolved decision state compared to the dynamic, evolving representations present during accumulation (Ofir & Landau, 2022, 2025). Third, the offset provides a time-locking point for analysis, which may reduce temporal variability and improve the ability to detect consistent neural responses. During the interval itself, the neural processes underlying temporal accumulation may unfold with variable latencies.

Within the post-offset period, we identified an early and a late component. Both were modulated by factors similar to those predicted for behavioral responses, suggesting that they track aspects of temporal decision-making. However, the functional roles of these components may differ. Across the literature, the late components show more consistent topography, are observed across different modalities, interval ranges, and tasks, and closely track context and behavioral biases (Baykan et al., 2023; Bueno & Cravo, 2021; Ofir & Landau, 2022; Özoğlu & Thomaschke, 2023). The early activity, although consistently found, is more task-dependent. For example, the topography of the effect seems to be modality-dependent, as intervals presented in different modalities (visual, somatosensory, or auditory) elicit an effect in sensors close to the respective sensory areas (Baykan et al., 2023; Bueno et al., 2024; Kruijne et al., 2020; Ofir & Landau, 2022; Özoğlu & Thomaschke, 2023). The variability in early activity may also reflect its closer ties to sensory processing. Although already modulated by temporal category, the early signal may retain modality-specific characteristics from the encoding stage. In contrast, the later signal may reflect a more abstract decision that has been transformed into categorical space, independent of the original sensory modality. This view is consistent with hybrid models of temporal perception that posit a transition of a local sensory-dependent pattern of activity to a shared representation of time (Merchant et al., 2013).

Timing models can be divided into two broad classes that make different predictions about how neural activity should change across different durations. Threshold-based models, such as SET (Gibbon, 1977), predict that the same accumulation process unfolds across intervals, with durations distinguished by the threshold levels they reach. Rate-based models, such as TopDDM (Balci & Simen, 2014; Simen et al., 2011), predict that neural activity rescales with duration, so that the same final pattern is reached at different speeds. Our results speak to both frameworks. We found that duration information was encoded along a distinct neural dimension, separate of categorical decisions, consistent with a dedicated accumulation signal as proposed by threshold-based accounts. We also observed that pre-offset activity systematically varied across reference intervals. This rescaling of pre-offset activity is considered a hallmark prediction of rate-based models (Balcı & Simen, 2016).

However, these findings do not straightforwardly adjudicate between the two models. Both models incorporate a decision stage in which the current interval is evaluated against a reference. This decision component should, by definition, scale with the ratio of the timed interval to the reference, regardless of the underlying timing mechanism. Our orthogonal design captured these categorical decision signals, which were present even during the pre-offset period, and evolved in parallel with duration encoding. Thus, the observed rescaling of activity could reflect changes in the decision variable, either in addition to or in place of changes in the timing mechanism. More broadly, our results suggest that claims about whether neural activity scales with duration must specify which component of temporal processing is being assessed. Without designs that dissociate timing from decision-making, apparent evidence for rescaling may conflate changes in the clock process with changes in the comparison or categorization process. The orthogonal design introduced here provides one such approach, enabling researchers to isolate timing-related neural dynamics from the decision processes that act upon them.

Regardless of whether the observed signals reflect the timing mechanism or the decision process, they were strongly modulated by recent experience. Classifier evidence was influenced by previous trials and, on equal trials, it was systematically predicted by the dynamically updated internal reference. This pattern indicates that the neural activity underlying temporal judgments is continuously shaped by contextual learning, as also observed in our behavioral findings. However, many of the present models treat timing and decision stages as operating with fixed parameters, assuming that learning has occurred before the test phase, even if for simplicity (Balci & Simen, 2014; Jazayeri & Shadlen, 2010; Machado et al., 2022). Our findings challenge this assumption, showing that learning can be captured in decoded neural signals. Models of temporal processing should incorporate mechanisms for updating rather than treating temporal judgments as comparisons against a stable standard.

Frameworks such as TopDDM (Balci & Simen, 2014; Balcı & Simen, 2016; Simen et al., 2011) are grounded in general decision-making theory rather than timing-specific mechanisms. This is an important difference, given that much of the timing literature has proposed dedicated models and neural signatures. However, dynamic updating of internal references is not unique to temporal tasks, and similar trial-by-trial adaptation has been reported across perceptual and cognitive domains (Lages & Treisman, 1998; Levari et al., 2018; Rigoli, 2019). Likewise, decision signals have been associated with evidence accumulation and linked to drift-diffusion frameworks in other domains (Luyckx et al., 2019; O’Connell et al., 2012; Summerfield et al., 2020; Twomey et al., 2015). This raises the question of how much of what we observe in temporal tasks reflects timing-specific computation versus general-purpose decision and learning mechanisms applied to temporal content. Future studies should compare neural signatures of temporal and non-temporal decisions to determine which aspects of the signals are unique to timing and which reflect domain-general processes.

## Acknowledgments

AMC was funded by the São Paulo Research Foundation (FAPESP), grant #2017/25161-8, CNPq grant #311721/2023-0, INCT NeuroComp #408389/2024-9 and INCT-ECCE, CNPq: #409051/2024-1, FAPESP: #2026/01540-9. MS was funded by the São Paulo Research Foundation (FAPESP) under grant #2016/24951-2. PMEC was funded by CNPq Universal, grant #409686/2021-2. This study was financed in part by the Coordenação de Aperfeiçoamento de Pessoal de Nível Superior – Brasil (CAPES) – Finance Code 001. The authors also wish to thank Mark Stokes for his contributions during the earlier stages of this project.

## Supplementary Figures

**Figure S01.**
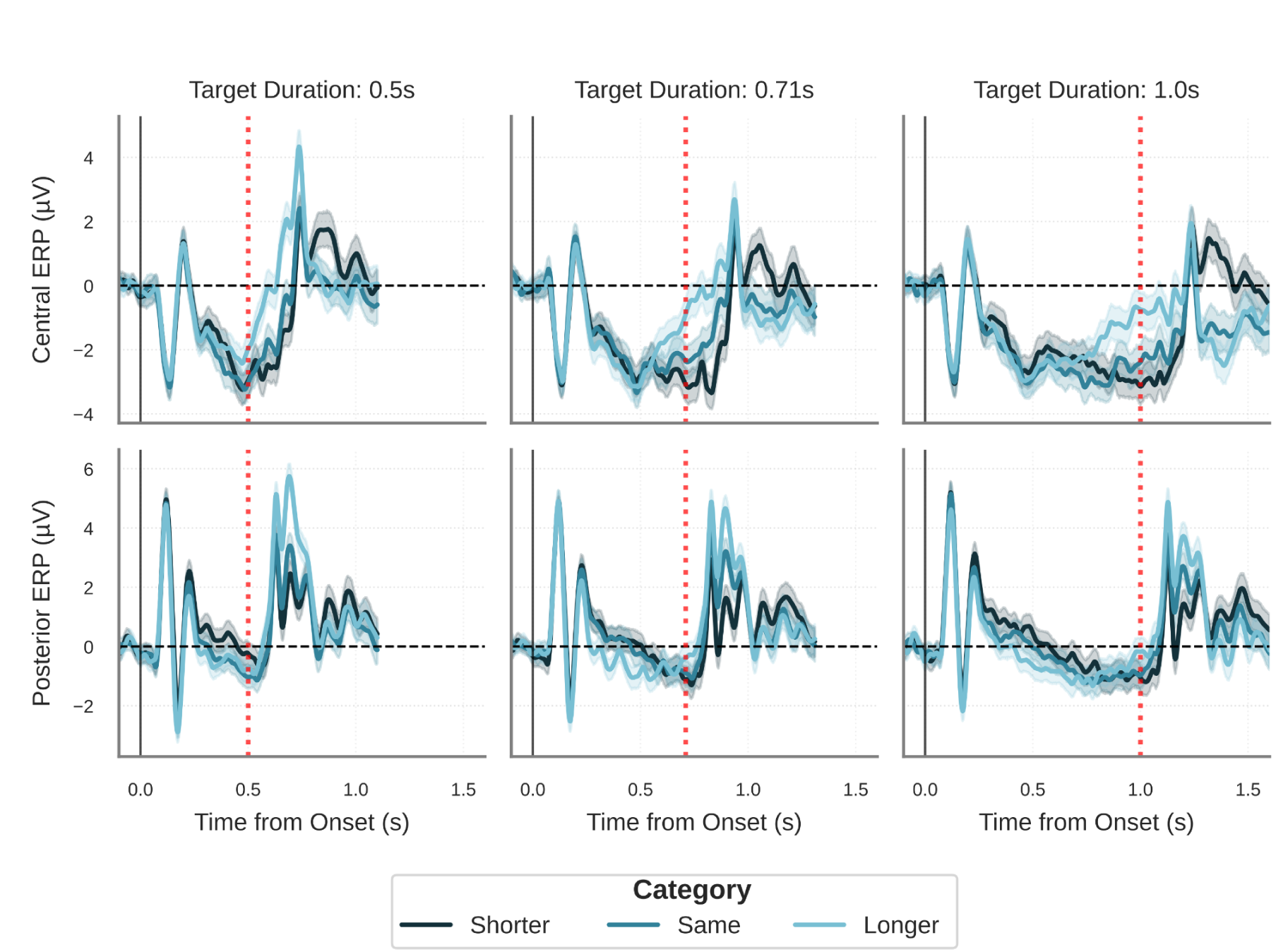
Event-related potentials in central and posterior ROIs as a function of target duration and temporal category. ERPs are shown for central (top row) and posterior (bottom row) regions of interest, separately for each target duration (0.5s, 0.71s, and 1.0s; columns). Data are aligned to stimulus <u>onset</u> (time 0, indicated by the solid black vertical line), with stimulus offset for each respective duration indicated by the red dotted vertical line. The three temporal categories (Shorter, Same, Longer) are shown in different shades. Lines represent the mean across participants, and shaded areas indicate 95% confidence intervals obtained via bootstrap resampling (n=1000).

